# Reinforcement learning modeling reveals a reward-history-dependent strategy underlying reversal learning in squirrel monkeys

**DOI:** 10.1101/2021.05.05.442852

**Authors:** Bilal A. Bari, Megan J. Moerke, Hank P. Jedema, Devin P. Effinger, Jeremiah Y. Cohen, Charles W. Bradberry

**Affiliations:** The Solomon H. Snyder Department of Neuroscience, Brain Science Institute, Kavli Neuroscience Discovery Institute, Johns Hopkins University, Baltimore, MD; NIDA Intramural Research Program, 251 Bayview Blvd, Suite 200, Baltimore, MD 21224, USA; Department of Pharmacology, University of North Carolina at Chapel Hill, Chapel Hill, NC

**Keywords:** reversal learning, reinforcement learning, squirrel monkeys, decision making, behavioral modeling

## Abstract

Insight into psychiatric disease and development of therapeutics relies on behavioral tasks that study similar cognitive constructs in multiple species. The reversal learning task is one popular paradigm that probes flexible behavior, aberrations of which are thought to be important in a number of disease states. Despite widespread use, there is a need for a high-throughput primate model that can bridge the genetic, anatomic, and behavioral gap between rodents and humans. Here, we trained squirrel monkeys, a promising preclinical model, on an image-guided deterministic reversal learning task. We found that squirrel monkeys exhibited two key hallmarks of behavior found in other species: integration of reward history over many trials and a side-specific bias. We adapted a reinforcement learning model and demonstrated that it could simulate monkey-like behavior, capture training-related trajectories, and provide insight into the strategies animals employed. These results validate squirrel monkeys as a model in which to study behavioral flexibility.

## Introduction

Psychiatry is in need of fundamental insights both so we may understand the psychological and neural basis of disease and so we can develop novel therapeutics. Comparative neuroscience is one approach that has historically led to the discovery of safe and effective pharmaceuticals (Markou et al., 2009). One common procedure is to combine animal models of psychiatric disease with commonly-used behavioral tasks, such as the forced swim test or elevated plus maze. Compounds can then be tested for their ability to ameliorate modeled symptoms (Kaiser and Feng, 2015; Flint and Shifman, 2008; Crawley, 2008; Dawson and Tricklebank, 1995; Nestler et al., 2002). While this approach has been fruitful, it has failed to fundamentally change our understanding about disease or lead to the discovery of truly novel therapeutics (Pangalos et al., 2007; Fenton et al., 2003). This is partly because preclinical behavioral models represent a major bottleneck in drug development (Tallman, 1999). Since psychiatric diseases affect higher-order cognitive processes, designing tasks that translate drug effects from animal models to humans is nontrivial. The standard library of tasks were designed to probe intuitive ideas about observable symptoms, not quantitative theories of cognitive processes.

A promising approach is to design behavioral tasks that probe the same psychological phenomena across species (Pike et al., 2021). These tasks may use different stimuli and motor responses, but attempt to isolate the same neurocomputational mechanisms. Theories about the relevant cognitive processes are used to form and test explicit hypotheses, in the form of models, about behavioral strategies (Wilson and Collins, 2019; Daw et al., 2011; Heathcote et al., 2015). From the perspective of computational psychiatry, these models in turn allow us to understand and quantify aberrant information processing in disease (Huys et al., 2016; Aylward et al., 2019; Gershman and Lai, 2020; Mason et al., 2017; Redish, 2004; Radulescu and Niv, 2019), as well as effects of therapy (Michely et al., 2020; Frank et al., 2007; Paulus et al., 2016).

Reversal learning is one popular task that is amenable to theory-based computational modeling (Behrens et al., 2007; Soltani and Izquierdo, 2019). In common variants of this task, subjects are presented with a choice between two stimuli, one associated with a high-value outcome (e.g. high probability or large volume of reward) and the other associated with a low-value outcome. Subjects begin the task with no knowledge about which stimulus is the better option. On each trial, subjects select one stimulus, receive the associated outcome, and repeat this process. Through trial-and-error, subjects learn the values of each stimulus. After some time, the two stimuli reverse in association (hence, reversal learning), so that the previously low-value stimulus becomes the high-value stimulus and vice versa. Importantly, these reversals are not cued, necessitating continual trial-by-trial learning to maximize reward. This task design is thought to engage mechanisms of flexible and rapid learning, impairments of which are implicated in a wide range of psychiatric disease (Swainson et al., 2000; Huys et al., 2013; Aylward et al., 2019; Remijnse et al., 2006; Brigman et al., 2009; Leeson et al., 2009; Izquierdo and Jentsch, 2012), including addiction (Porter et al., 2011; Ersche et al., 2011).

Although behavior on these tasks is typically reported using simple summary statistics (average performance, trials to reach a criterion, etc.), richer insight can be gleaned with reinforcement learning modeling. Reinforcement learning is a framework that formalizes learning from environmental feedback (Sutton and Barto, 1998), and has provided a number of tractable algorithms that have delineated numerous structure-function relationships in the nervous system (Schultz et al., 1997; Samejima et al., 2005; Bari et al., 2019; Ottenheimer et al., 2020; Grossman et al., 2020; O’Doherty et al., 2004). Among the most commonly-applied algorithms are the class that iteratively learn stimulus values over many trials and choose based on the relative values of the stimuli. Reinforcement learning models of reversal learning have been used to explain behavioral data in species as diverse as rodents (Harris et al., 2020; Metha et al., 2020), macaques (Costa et al., 2016), and humans (Kanen et al., 2019). Importantly, reinforcement learning models are generative models — that is, they are capable of simulating behavior, a premise which we capitalize on in this manuscript.

Here, we trained squirrel monkeys on a deterministic image-based reversal learning task. Squirrel monkeys are New World primates widely used in biomedical research, primarily due to their small size (<1kg), ease of handling, and adaptation to laboratory conditions (Abee, 2000). From the perspective of comparative neuroscience, squirrel monkeys help span the massive genetic, anatomical, and behavioral gap between rodents and humans (Boinski, 1999). They may also prove to be a useful preclinical model for development of optogenetic-based interventions (O’Shea et al., 2018).

Our objective was to determine if squirrel monkeys solve reversal learning tasks using a strategy compatible with trial-by-trial reinforcement learning, and to isolate parameters of cognitive flexibility to employ in future studies. First, we demonstrate that squirrel monkeys do not adopt the optimal win-stay/lose-shift strategy required to optimize reward accumulation in this task but rather integrate reward over many trials. We fit a number of reinforcement learning models and found that a standard Rescorla-Wagner model fit best, similar to reversal learning models in other species. We show that this model simulates realistic behavior, providing a convincing platform for making inferences about behavioral strategy. Finally, we use the recovered parameters to define how the behavioral strategy develops with training.

## Methods

### Subjects

A total of 13 (9 of which met behavioral criteria) adult male squirrel monkeys (*Saimiri sciureus*) with less than 1 year of training on behavioral touchscreen tasks were housed individually under controlled temperature and humidity on a 12/12-h light-dark cycle (lights on from 0700 to 1900h). Monkeys weighed 867-1113 g (mean: 965 g) and were maintained on a diet of primate chow (LabDiet High Protein Monkey Biscuits; PMI Feeds, St. Louis, MO) with continuous access to water in the home chamber. Environmental enrichment, including fresh fruits and vegetables, was provided on a daily basis. The maintenance and experimental use of animals was carried out in accordance with the 2011 Guide for Care and Use of Laboratory Animals. All experimental protocols were approved by the Animal Care and Use Committee of the National Institute on Drug Abuse Intramural Research Program.

### Apparatus

Experiments were conducted in sound-attenuating chambers equipped with a 15” touchscreen (Elo TouchSystems, Menlo Park, CA), mounted in a panel 14.25” from the floor of the chamber. Centered 1.5” below the touchscreen and extending 2” into the chamber was a well into which measured volumes of 30% sweetened condensed milk (Eagle Foods, Richfield, OH) could be delivered through a line connected to a syringe pump (Harvard Apparatus, South Natick, MA) located outside the chamber. Monkeys were seated in custom-built acrylic chairs facing the touchscreen panel. A computer and software program (E-Prime Professional 3.0; Psychology Software Tools, Inc., Sharpsburg, PA) controlled the parameters of the experimental program and data collection.

### Behavioral task and training

Prior to introducing the reversal task, monkeys were trained on a task in which they chose between different quantities of milk (0.075-0.3 ml/kg) represented by unique stimuli on the touchscreen. Monkeys registered a choice by physically touching the display for 100 ms; this was paired with a tone and allocation of the associated reward into the well below the touchscreen. A house light and speakers inside the experimental chambers provided illumination and white noise. Training sessions were generally carried out five days a week (Monday-Friday) and lasted 30-60 min.

Following this training task, monkeys conducted an image-based deterministic reversal learning task. These sessions began with a Discrimination block, in which monkeys were presented with two novel images selected randomly from a large library of images. One image was associated with a big reward (large volume of milk; the ‘correct’ choice) and the other image was associated with a small reward (small volume of milk; the ‘incorrect’ choice). Monkeys registered a choice by physically touching one of the visual stimuli on the display and received the associated reward from a reward port. Following an 8-12 second intertrial interval, the images were presented again on the next trial, with left/right positions randomized between trials. Monkeys made choices until they reached a performance threshold of 80% correct in the past 15 trials, after a minimum 20 trial block length. Once this threshold was reached, a Reversal block was initiated, in which the two image associations reversed so the image previously associated with big reward was now associated with small reward, and vice versa for the other image. Monkeys again performed until they reached the performance threshold, at which point a new Discrimination block was initiated and two new images were randomly sampled from the library. Monkeys typically performed for 150 trials, although some sessions were shorter due to reduced motivation. The large reward (0.13-0.24 ml/kg) was four times larger than the small reward (0.03-0.06 ml/kg).

### Data analysis

All 13 monkeys completed at least 60 sessions and at least 1 block per session on average. Monkeys that reached an average performance threshold of 54% across all reward blocks and all sessions were included, yielding 9 monkeys in the final dataset. Monkeys performed an average of 121 sessions (range 66-135).

All choices that yielded big (small) reward were labeled as correct (incorrect). Performance was defined as the fraction of correct choices in a session. To generate reward history regressions, we arbitrarily coded one image as “image 0” and the other image as “image 1” for each set of presented images and fit the following random effects logistic regression

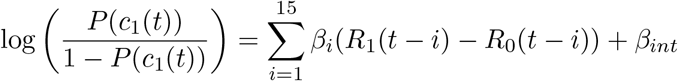

where *c*_1_(*t*) = 1 for a choice to “image 1” and 0 for a choice to “image 0”. *R*(*t*) = 1 if big reward was delivered for that image on trial *t* and 0 otherwise. We included monkey-level and session-level (nested within monkey) random effects for the intercept.

We generated errorbars for performance within blocks (Figures 2D, 4D) by computing bootstrapped 95% confidence intervals from 1,000 bootstrap samples of the mean.

In generating image-based win-stay/lose-shift and mutual information metrics, we excluded the first trial of each Discrimination block, as new images were presented on these trials. The mutual information between stay/shift (to image) and reward on the previous trial was calculated as

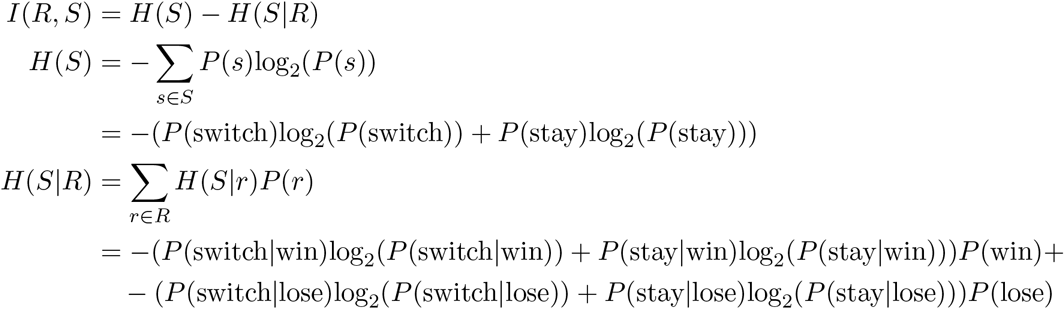

where *I*(*R, S*) is the mutual information, *S* = {switch,stay} on the current trial, and *R* = {win,lose} on the previous trial.

Side bias was defined as 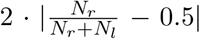 where *N_r_* and *N_l_* are the total rightward and leftward choices in a session, respectively. Side bias = 0 if there are an equal number of leftward/rightward choices and 1 if all choices are exclusively to one side. The entropy of the side chosen distribution was calculated as

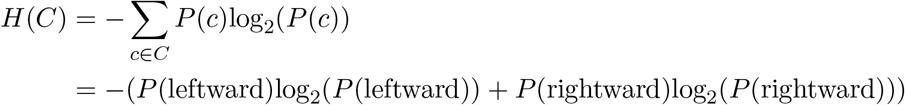

where *C* = {leftward,rightward}.

All regressions relating behavioral metrics to sessions number were random effects linear regressions with monkey-level random effects for slope and intercept.

### Reinforcement learning models

We developed a number of reinforcement learning models based on the Rescora-Wagner model. Our chosen model took the following form for updating image values

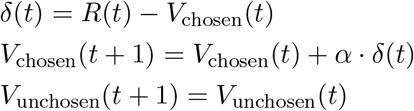

where values were initialized with *V_i_* = 0 at the beginning of each Discrimination block. In this model, the chosen image’s value is updated based on the discrepancy between prediction and reward (reward prediction error, *δ*(*t*)). The unchosen image’s value remains unchanged. Image values were fed into a softmax function to generate choices according to

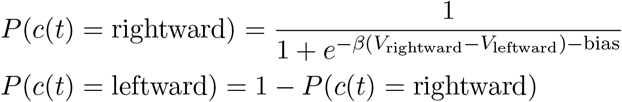

We fit a number of model variants. First, we considered a set of noise models testing whether behavior could be explained as random, biased, perseverative (1-trial-back choice autocorrelation), or biased + perseverative. Within the space of Rescorla-Wagner models, we considered models with all possible combinations of the following: one learning rate, two learning rates (separate learning of positive and negative reward prediction errors), forgetting of unchosen image values to 0, rewards coded as [0.25 1] (since the small reward was 25% the volume of the large reward; only for models with forgetting), and learning of action values (i.e. learning values for left-ward/rightward actions). Each of these models also included permutations for nuisance parameters (none, side bias, perseveration, or side bias + perseveration). We additionally considered a set of models that augmented each model to allow for a mixture of image-based win-stay/lose-shift and reinforcement learning. We considered one final model (a variant of our chosen model) that explicitly accounted for a reversal mechanism. In this model, reward prediction errors symmetrically updated image values - if one image value was increased by *δ*(*t*), the other image value was decreased by *δ*(*t*), while bounding image values by [0 1]. In all models reward values were coded as 0 and 1 for small reward and big reward, respectively (except for the variant where they were coded as [0.25 1]). In total, we considered 105 model variants.

We developed a metric, the maximal trial-by-trial change in P(choice), to capture the interaction between the learning rate, *α*, and the inverse temperature, *β* (Figure 6F,J, S6C). For a given *α* and *β*, we assumed the largest reward prediction error, *δ*(*t*) = 1. This yields *V*_chosen_(*t* + 1) = *V*_chosen_(*t*) + *α* · *δ*(*t*), which can be simplified as *V*_chosen_(*t* + 1) − *V*_chosen_(*t*) = *α*. In other words, the value function is increased by *α* in response to the largest possible reward prediction error in this task. We then calculated the change in P(choice) around the inflection point of the softmax function (at P(choice) = 0.5) since the slope is steepest at this point. This yields the maximum trial-by-trial change in P(choice).

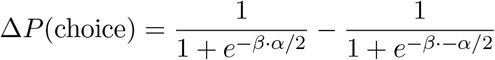

### Model fitting

Models were fit to individual session data with maximum likelihood estimation, with 10 starting points to avoid finding local minima. To determine which models fit the data most parsimoniously, we used the Bayesian information criterion, which penalizes models with additional parameters. The above reinforcement learning model fit best for the most monkeys (Table S1).

This model is notable for several reasons. First, none of the noise models fit best for the monkeys included in this dataset. Because Bayesian information criterion, relative to other metrics (like Akaike information criterion), favors simpler models, this suggests that Rescorla-Wagner-like learning is a key feature of behavior. Second, although reinforcement learning models with two learning rates are often fit to animal and human data, we found that none of the models with two learning rates were selected. Third, none of the win-stay/lose-shift models provided better fits than the complementary model variants without win-stay/lose-shift. This includes four win-stay/lose-shift models that did not include Rescorla-Wagner-style learning (a mixture of the four noise models + win-stay/lose-shift).

### Model recovery

For the best model, we took the parameter estimates for each session and generated fictive data according to the same model. We then fit all 105 models to this synthetic dataset and found that the true generative model was selected for 9 of the 9 simulated monkeys. This shows that our model recovery procedure could indeed recover our chosen model.

We also conducted a parameter recovery exercise with these models and found that the difference between actual and recovered parameters had a mode of 0 for all three parameters.

### Hierarchical Bayesian model fitting

To obtain partially-pooled parameter estimates (i.e., less noisy estimates, especially since *α* and *β* tend to compensate for one another (Daw et al., 2011; Ballard and McClure, 2019)), we refit the best reinforcement learning model using a hierarchical Bayesian framework. We used MATLAB (Mathworks), the probabilistic programing language Stan (https://mc-stan.org), and the MATLAB interface, MatlabStan (https://mc-stan.org/users/interfaces/matlab-stan). We constructed hierarchical models separately for each monkey, with monkey-level parameters to govern session-level parameters for learning. Priors over monkey-level parameters, from which session-level means were drawn, were set as

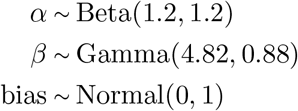

where the priors for *α* and *β* were taken from the literature (Kanen et al., 2019; Den Ouden et al., 2013; Gershman, 2016). The gamma distribution was parameterized in terms of shape and scale. For all monkey-level variances, we used Cauchy^+^(0, 1). For session-level *α* and *β*, we again used beta and gamma distributions, reparameterized in terms of mean and variance with parameters drawn from monkey-level distributions. Session-level bias was normally distributed, with mean and variance drawn from monkey-level distributions. Parameter values reported in Figure 6 are the means of the session-level posteriors. Distributions in Figure 6A-C are posteriors over monkey-level means.

## Results

### Reward history and side bias inform strategy

We developed a deterministic reversal learning task in which chaired squirrel monkeys chose between two simultaneously presented images for delivery of milk reward. Images were presented in blocks of trials, and in each block, one image was associated with big reward and the other was associated with small reward. On each trial, monkeys were presented with two images, each on the left/right half of a touchscreen and physically touching an image yielded reward (Figure 1A). Selecting the big reward image (which we call the correct choice) for 80% of the past 15 trials triggered a block transition, uncued to the monkey. Blocks switched between Discrimination blocks and Reversal blocks (Figure 1B). At the beginning of each Discrimination block, two images were randomly selected from a large library of images and assigned to big/small reward. At the beginning of each Reversal block, the two images swapped reward contingencies. On average, sessions lasted for 146 (SD 14) trials and monkeys completed 4.8 (SD 1.9) blocks.

**Figure 1:**
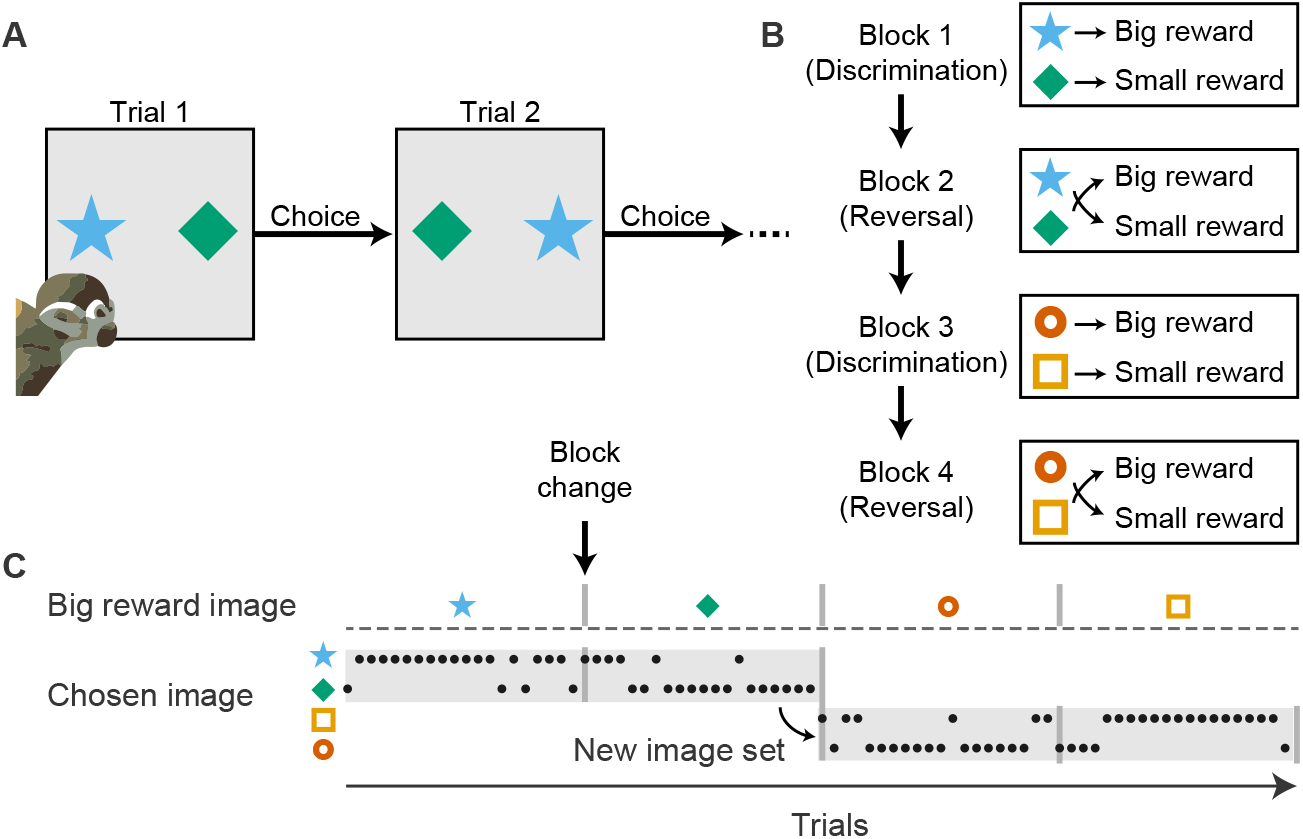
Reversal learning task design. (A) Squirrel monkeys chose between two images presented on the left and right halves of a touchscreen. A choice was registered by physically touching either visual stimulus on the display. One image was deterministically associated with big milk reward, and the other image was associated with small milk reward. Image locations were randomly displayed on the left and right halves of the screen on separate trials. (B) Monkeys performed sequences of Discrimination and Reversal blocks. At the beginning of each Discrimination block, two new images were randomly sampled from a large library of images and each image was randomly assigned to big or small reward. At the beginning of each Reversal block, the two images switched reward contingencies. Block transitions were triggered by a threshold of 80% correct responses (response to the big reward image) in the past 15 trials, after a minimum of 20 trials. These transitions were unsignaled, requiring the animal to use reward feedback to guide decisions. (C) Example choice behavior demonstrates the flexibility of behavior at Discrimination → Reversal and Reversal → Discrimination block transitions.

The optimal strategy in this task is an image-based win-stay/lose-shift policy: select the same image if it yielded big reward on the previous trial, switch if it yielded small reward. After training, monkeys reliably switched their choices at block transitions, when contingencies switched (Figure 1C, S1A). However, although animals performed significantly better than chance (Wilcoxon signed-rank test, p < 0.01), they performed worse than win-stay/lose-shift, a optimal strategy that would yield the large reward on ~ 96% of trials (Figure 2A). To understand how monkeys solved this task, we fit logistic regression models to predict choice as a function of reward history. Unlike the optimal policy, which maintains a memory of reward on just the most recent trial, monkeys maintained a recency-weighted memory of reward history up to 10 trials in the past to inform choices (Figure 2B). We also found that monkeys were faster to transition from Reversal → Discrimination blocks than from Discrimination → Reversal blocks (Figure 2C,D). A two-way ANOVA (Block Type x Trials) was performed separately for trials before and after the block transition. Before the block transition, there was no significant effect of Block Type (*F*_1,96_ = 1.32, p = 0.25). After the transition, this effect became significant (*F*_1,240_ = 270.13, p < 0.0001). Performance in Discrimination blocks was better than in Reversal blocks (mean [95%CI], 0.685 [0.679 – 0.690] fraction correct in Discrimination blocks, 0.576 [0.568 – 0.583] fraction correct in Reversal blocks). This asymmetry in block performance is consistent with the idea that Reversal blocks, but not Discrimination blocks, require unlearning of previously-learned associations in addition to learning new associations.

**Figure 2:**
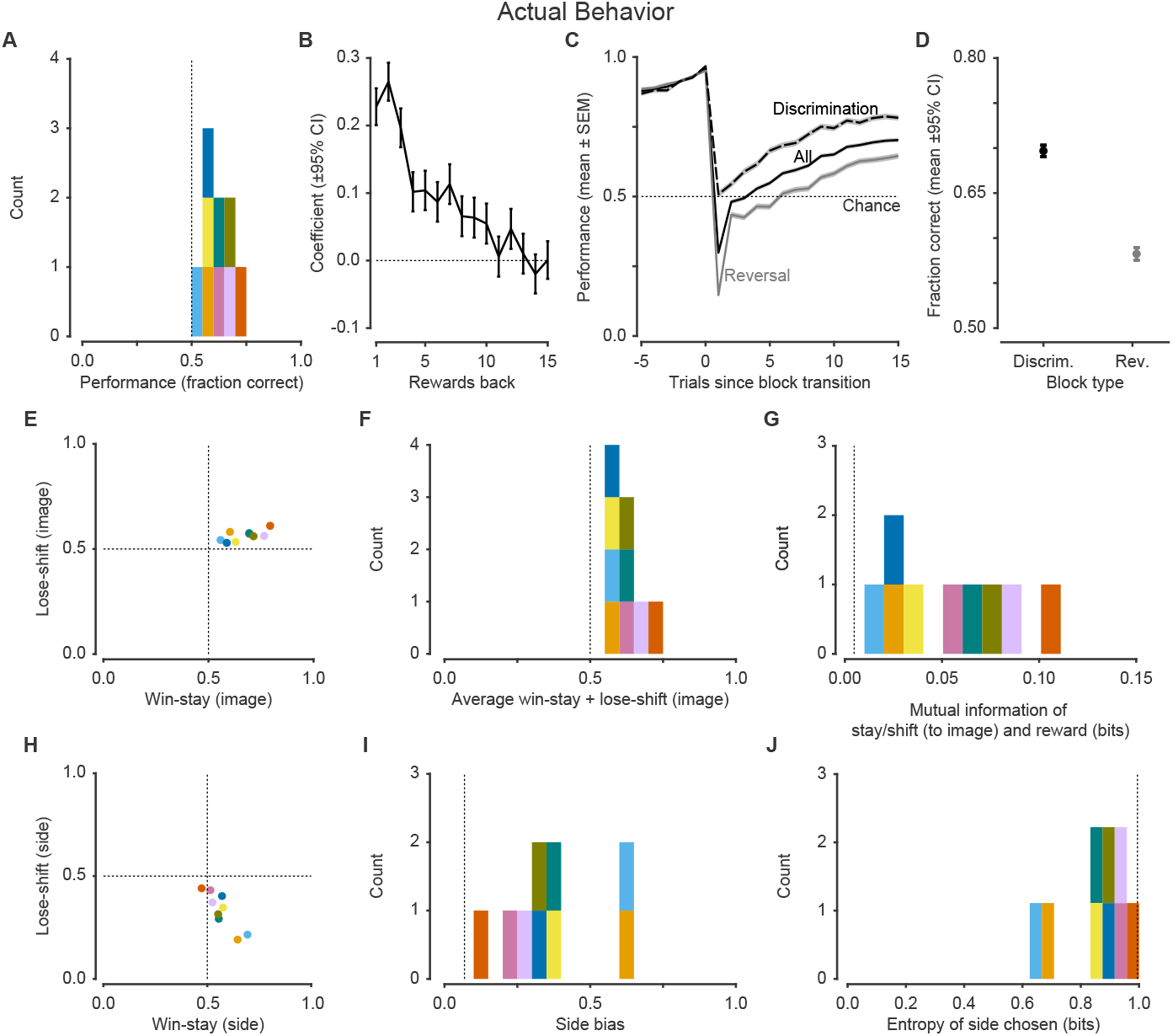
Behavioral features demonstrate reward sensitivity and side bias. (A) Performance was significantly better than chance (50%, dashed line). (B) Logistic regression coefficients for choice as a function of reward history. (C) Performance at block transitions for all blocks, and separately for Discrimination and Reversal blocks. Relative to Reversal blocks, monkeys were faster to improve performance during new Discrimination blocks. The increase in performance prior to block transitions is because transitions were triggered by good performance. (D) Performance was better in Discrimination blocks relative to Reversal blocks. (E) Image-based win-stay and lose-shift were both greater than 0.5, demonstrating that animals learned from both wins (big reward) and losses (small reward) to guide decisions. (F) The average win-stay + lose-shift, which can be taken as a proxy for the strength of reward-guided behavior, was greater than 0.5 (dashed line). Values close to 0.5 are consistent with reward-insensitive behavior and values of 1.0 are consistent with a perfect win-stay lose-shift strategy. (G) The mutual information between stay/switch and reward on the previous trial. Mutual information quantifies how much better we can predict the strategy (stay vs switch) if we know the reward received on the previous trial (dashed line is from simulated random behavior). (H) Side-based win-stay and lose-shift highlight a side bias, where animals largely stay. (I) Side bias, which is 1 if choices are exclusively to one side and 0 if they are uniformly split, was widely distributed (dashed line is from simulated non-side-biased behavior). (J) The entropy of the side chosen distribution showed a similarly wide distribution. Entropy of 1 indicates choices were uniformly split and entropy of 0 indicates choices were exclusively to one option (dashed line is from simulated non-side-biased behavior). Colors denote individual monkeys and are consistent between figures.

Following reward, monkeys can implement two distinctive strategies: choose to repeat choices to the same image, regardless of side (image-based win-stay) or repeat choices to the same side, regardless of image (side-based win-stay). To better define monkeys’ behavioral strategy, we first quantified win-stay and lose-shift tendencies in image-based coordinates (Figure 2E). Each point indicates the fraction of trials in which monkeys stayed after receiving big reward (*x*-axis) and shifted after receiving the small reward (*y*-axis). Plotted this way, points in the top-right and bottom-left quadrants indicate reward-sensitive behavior (win-stay/lose-shift and win-shift/losestay, respectively) and points in the top-left and bottom-right quadrants indicate reward-insensitive behavior (shift and stay, respectively, regardless of reward). All monkeys were in the top-right quadrant, indicating that they were demonstrating both win-stay and lose-shift behavior following outcomes (Wilcoxon signed-rank test, p < 0.01 for each). We quantified reward sensitivity by computing the average win-stay + lose-shift per animal. This metric is 1.0 for perfect win-stay/lose-shift behavior, 0.5 for reward-insensitive behavior, and intermediate for reward-sensitive behavior. Consistent with prior analyses, animals demonstrated reward-sensitive behavior (Wilcoxon signed-rank test, p < 0.01). However, one shortcoming of this metric is it places equal emphasis on win-stay and lose-shift. Because *P*(lose) is fairly low in this task, behavior in response to losses does not impact overall performance as strongly as response to reward. To work around this pitfall, we computed the mutual information between reward on the previous trial and stay/switch behavior on the current trial, as it accounts for the base rate of *P*(lose) (Figure 2G). Perfect win-stay/lose-shift behavior results in 1 bit of information and reward-insensitive behavior results in 0 bits. Consistent with the average win-stay + lose-shift analysis, monkeys demonstrated reward-sensitive behavior (Wilcoxon signed-rank test, p < 0.01).

One notable behavioral suboptimality we observed was a bias towards a particular side (a rightward side bias can be seen in Figure S1A) (Friedman et al., 2017; Bari et al., 2019). To better understand this side bias, we quantified win-stay/lose-shift in side-based coordinates (Figure 2H). Most monkeys fell in the bottom-right quadrant, consistent with a reward-insensitive tendency to favor a particular side (win-stay > 0.5 and lose-switch < 0.5, Wilcoxon signed-rank test, p < 0.01 for each). We quantified side bias with a side bias metric (0 for uniformly split choices, 1 for exclusive choice of one side) and observed a wide distribution, indicating an average tendency for side biased behavior (Figure 2I; Wilcoxon signed-rank test, p < 0.01). The entropy of the side chosen distribution was similarly wide (1 bit for uniformly split choices, 0 bits for exclusive choice of one side; Figure 2J; Wilcoxon signed-rank test, p < 0.01)).

Taken together, these results argue that monkeys solve this task by integrating reward over many trials to inform choices, and that this strategy is corrupted by a side bias.

### Monkeys develop reward sensitivity and reduce side bias with training

The wide range of performance allowed us to relate behavioral performance to the behavioral metrics we defined. First, we found that the average win-stay + lose-shift increased with better performance (linear slope 0.82, *t*_7_ = 28.23, p < 0.0001), consistent with the optimality of image-based win-stay/lose-shift (Figure 3A). Similarly, the mutual information between reward and stay/shift increased with performance (Figure 3B; linear slope 0.50, *t*_7_ = 19.54, p < 0.0001). Next, we focused on the side bias (Figure 3C). We found that for poor performance, side bias was generally high and decreased with improved performance (Mean: linear slope −1.98, *t*_7_ = −3.17, p = 0.016). The entropy of the side chosen distribution increased with improved performance (Figure 3D; Mean: linear slope 1.31, *t*_7_ = 2.84, p = 0.025).

**Figure 3:**
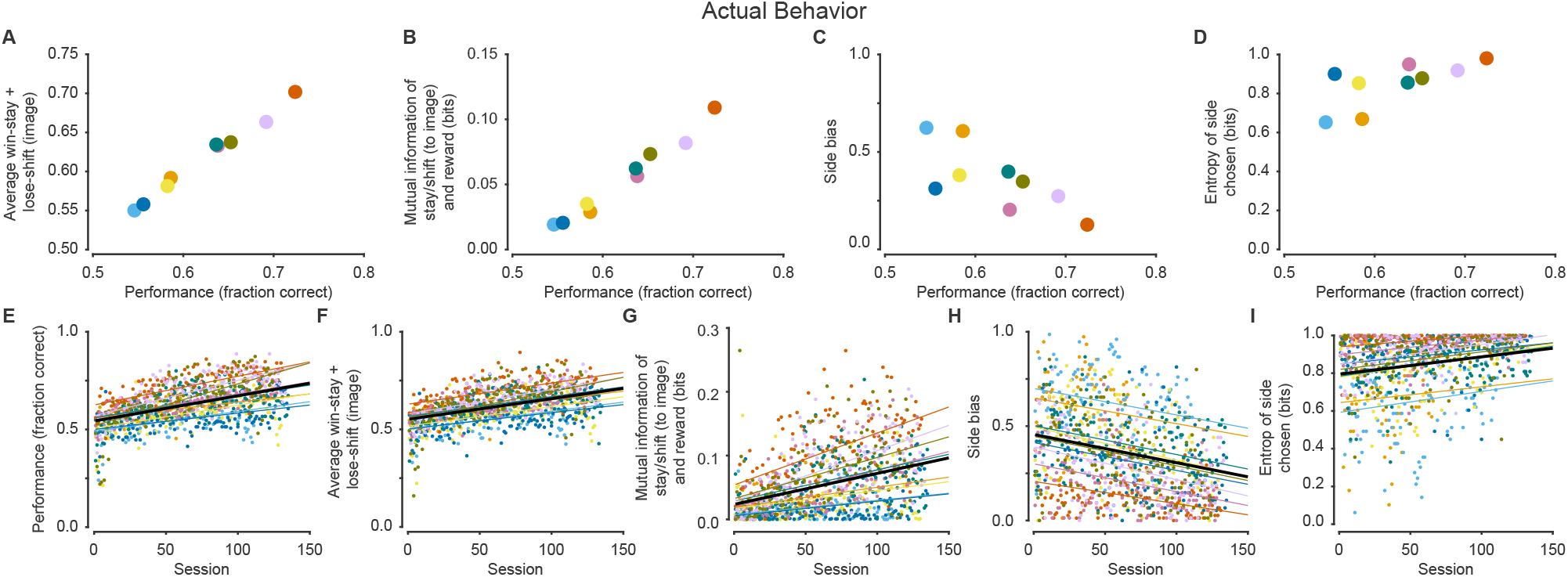
Relationship between performance, reward sensitivity, side bias, and training. (A) The average win-stay + lose-shift increased with increased performance. (B) The mutual information between stay/shift and reward increased with performance > 0.5. (C) Side bias was higher when performance was closer to 0.5 and reduced when performance was better. (D) Similarly, the entropy of the side chosen distribution increased (i.e. left/right choices become more random) when performance improved. (E) Performance improved with more sessions performed. (F) The average win-stay + lose-shift improved with training. (G) The mutual information between stay/switch and reward increased with training. (H) Side bias decreased with training. (I) Similarly, the entropy of the side chosen distribution increased with training. Black line shows the fixed effect and thin colored lines show individual monkey random effects. Colors denote individual monkeys and are consistent between figures.

The large number of sessions per monkey additionally allowed us to quantify the effects of training. First, we found that performance increased with the number of sessions (Figure 3E; linear slope 1.3 × 10^−3^, *t*_1091_ = 7.85, p < 0.0001). This was partly due to increased reward-sensitivity to images. The average win-stay + lose-shift increased with training (Figure 3F; linear slope 1.1 × 10^−3^, *t*_1091_ = 11.04, p < 0.0001). Similarly, the mutual information between reward and stay/switch increased (Figure 3G; linear slope 4.86 × 10^−4^, *t*_1091_ = 5.58, p < 0.0001). The improvement in performance was also partly due to a decrease in side bias. The side bias decreased with training (Figure 3H; linear slope −1.5 × 10^−3^, *t*_1091_ = −8.51, p < 0.0001). Similarly, the entropy of the side chosen distribution increased with training (Figure 3I; linear slope 9.17 *× −*4, *t*_1091_ = 6.64, p < 0.0001). In summary, monkeys improve with training, partly due to increased reward sensitivity to images, and partly due to a decrease in side bias.

### Reinforcement learning modeling captures key features of behavior

Since we found that monkeys integrated rewards over many trials, we adapted the Rescorla-Wagner model, a commonly-used model in reinforcement learning (Rescorla, 1972). This model maintains a running estimate of the values of images and chooses based on the relative values of the presented images. Image values are learned by recency-weighted reward history, which allows the model to adapt behavior flexibly when reward contingencies change. We considered a number of model variants: equivalent vs differential learning from better-than-expected and worse-than-expected outcomes, forgetting of unchosen image values, learning the values of actions (e.g. if leftward choices were recently rewarded, then increase probability of leftward choices), mixtures of reinforcement learning and win-stay/lose-shift strategies, and nuisance parameters, like side bias and choice autocorrelation. We fit individual sessions using maximum likelihood estimation and selected the best model using Bayesian information criteria, which selects the best-fit model while penalizing overly complex models. The best model was among the simplest — learning of image values with equivalent learning from better/worse outcomes and a side bias mechanism (Table S1). Importantly, this model was strongly preferred over noise models (which include nuisance parameters but no learning of image values), suggesting that learning image values was consistent with real behavior. Armed with a simple and tractable model, we sought to determine how well it described real behavior.

First, we observed that the model fit behavioral data well (data not shown). However, model fits run the risk of overfitting to data (Palminteri et al., 2017). A stronger approach is to take advantage of the generative modeling framework: simulate fictive data, and assess how well simulated data matches real behavioral data. Visually, we observed a correspondence between raw behavior and simulations (Figure S1B). Across all simulated monkeys, we observed that simulated behavior performed better than chance, similar to real behavior (Figure 4A; Wilcoxon signed-rank test, p < 0.01). Simulated behavior exhibited a dependence on reward history many trials into the past (Figure 4B). Like real monkeys, simulated monkeys were faster to transition to new Discrimination blocks than to new Reversal blocks (Figure 4C). There was no significant effect of Block Type prior to block transitions (*F*_1,96_ = 0.50, p= 0.48), which became significant after the transition (*F*_1,240_ = 274.78, p < 0.0001). Performance for Discrimination and Reversal blocks were comparable to real behavior (Figure 4D; Mean [95%CI], Discrimination: 0.685 [0.679 – 0.691], Reversal: 0.576 [0.568 – 0.584]).

**Figure 4:**
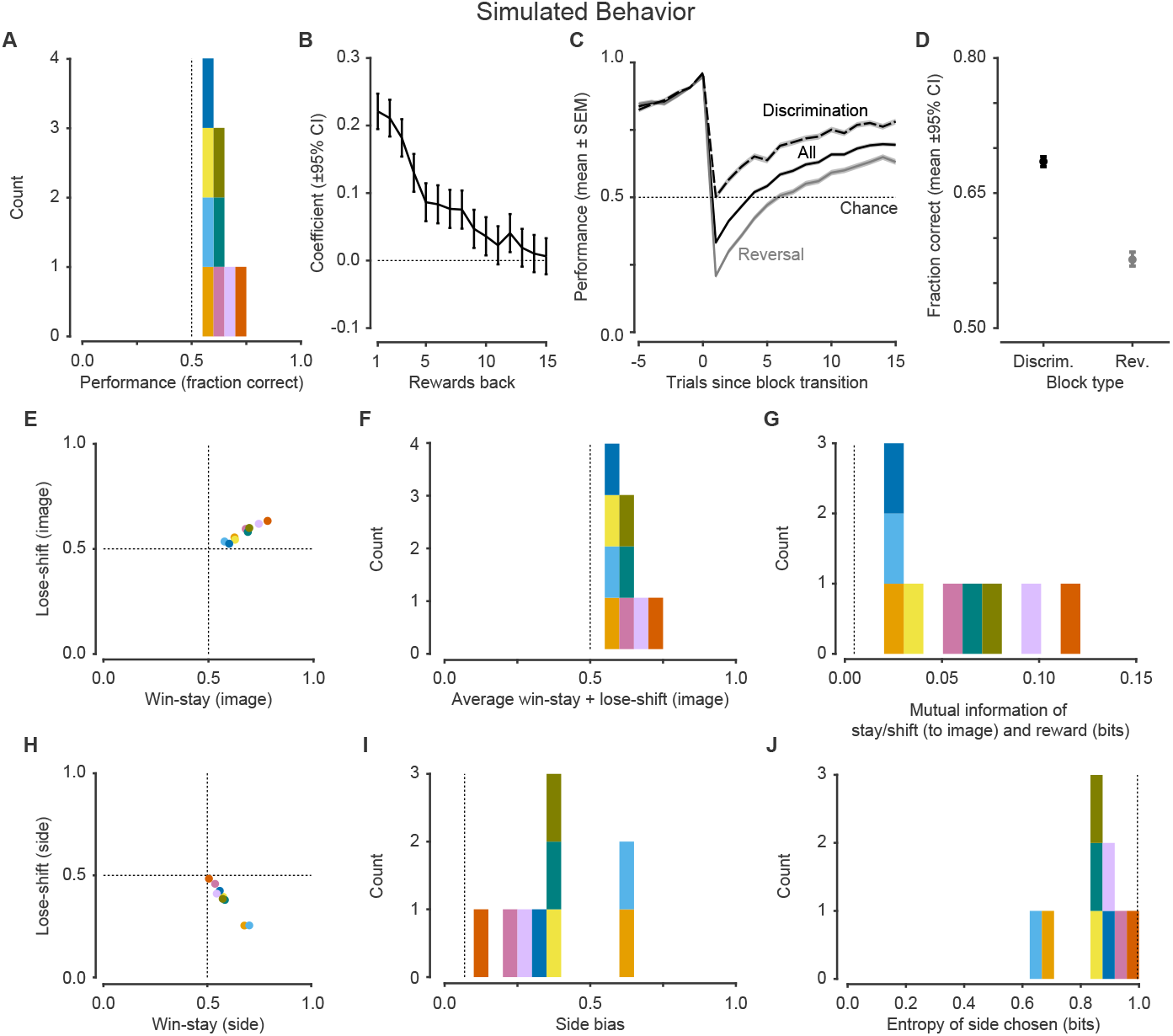
Simulated behavioral features demonstrate monkey-like reward sensitivity and side bias. (A) Distribution of performance was better than chance. (B) Logistic regression coefficients for choice as a function of reward history shows dependence for many trials in the past. (C) Performance at block transitions for all blocks, and separately for Discrimination and Reversal blocks. Like actual performance, pre-transition simulated data had no significant effect of Block Type which became significant after the transition. (D) Simulated performance was better in Discrimination blocks relative to Reversal blocks. (E) Image-based win-stay and lose-shift were both greater than 0.5. (F) The average win-stay + lose-shift was greater than 0.5. (G) The mutual information between stay/switch and reward on the previous trial is greater than random behavior. (H) Side-based win-stay and lose-shift demonstrates a side bias. (I) Side bias distribution. (J) The entropy of the side chosen distribution. Colors denote individual monkeys and are consistent between figures.

Simulated behavior exhibited features of reward-sensitivity to images, with image-based win-stay and lose-shift both > 0.5 (Figure 4E; Wilcoxon signed-rank test, p < 0.01 for each). The average win-stay + lose-shift was > 0.5, indicating reward-sensitive behavior (Wilcoxon signed-rank test, p < 0.01) and mutual information between reward and stay/switch was similarly skewed away from 0 bits, indicating reward sensitivity (Wilcoxon signed-rank test, p < 0.01; Figure 4F,G). Simulated behavior also exhibited suboptimal features of side bias (Figure 4H; win-stay > 0.5 and lose-switch < 0.5, p < 0.01 for each). Side bias and the entropy of the side chosen distributions were similarly wide (Figure 4I,J). On a monkey-by-monkey basis, there was a strong correspondence between each of these metrics for real and simulated data (Figure S2).

We addressed the relationship between simulated behavioral metrics and performance. We observed a strong linear dependence between performance and average win-stay + lose-shift (Figure 5A; linear slope 0.91, *t*_7_ = 47.85, p < 0.0001). There was likewise a strong association between performance and mutual information between reward and stay/switch (Figure 5B; linear slope 0.57, *t*_7_ = 23.63, p < 0.0001). Side bias decreased with performance (Figure 5C;linear slope −1.99, *t*_7_ = −3.06, p = 0.18). Entropy of the side chosen distribution similarly increased with performance (Figure 5D; linear slope 1.33, *t*_7_ = 2.68, p = 0.03).

**Figure 5:**
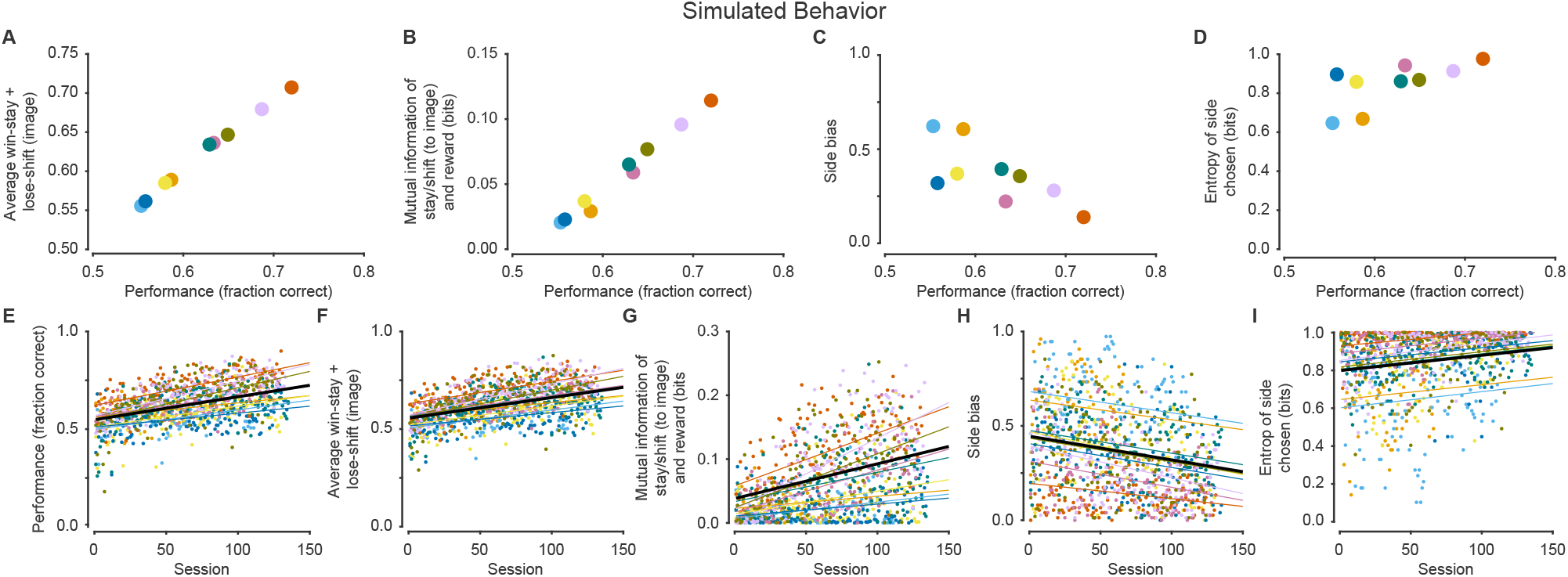
Simulations show similar relationships between performance, reward sensitivity, side bias, and training. (A) Average win-stay + lose-shift showed a positive relationship with performance. (B) Mutual information between stay/shift and reward as a function of performance. (C) Side bias as a function of performance. (D) Entropy of side chosen as a function of performance. (E) Performance improved with training. (F) Average win-stay + lose-shift improved with training. (G) Mutual information between stay/switch and reward increased with training. (H) Side bias decreased with training. (I) Entropy of the side chosen distribution increased with training. Black line shows the fixed effect and thin colored lines show individual monkey random effects. Colors depict individual monkeys and are consistent across figures

We also addressed the relationship between behavioral metrics and training. Simulated behavior exhibited an increase in performance with training (Figure 5E; linear slope 1.12×10^−3^, *t*_1091_ = 8.00, p < 0.0001). The average win-stay + lose-shift improved with training (Figure 5F; linear slope 1.05 × 10^−3^, *t*_1091_ = 8.56, p < 0.0001) and the mutual information between reward and stay/switch improved with training (Figure 5G; linear slope 5.44 × 10^−4^, *t*_1091_ = 4.65, p < 0.0001). Side bias decreased with training (Figure 5H; linear slope −1.25 × 10^−3^, *t*_1091_ = −6.35, p < 0.0001) and the entropy of the side chosen distribution similarly increased (Figure 5I; linear slope 8.10 × −4, *t*_1091_ = 6.90, p < 0.0001).

Taken together, these results argue that our reinforcement learning model with two core features — learning of image values, and a side bias — is sufficient to capture key features of real behavior.

### Model parameters provide interpretable insight into behavioral strategy

A key advantage of our generative modeling approach, beyond traditional summary statistics (e.g. mean performance, block lengths, etc), is the ability to provide intuitive explanations for how behavior was generated. Our model has two key components - a learning component and a decision component. The learning component determines the image values and the decision component turns the relative image values into a decision. The model has three parameters, which we detail below: learning rate (*α*), inverse temperature (*β*), and side bias (Figure S3A).

The learning rate, which affects the learning component, determines how quickly image values are updated following an outcome (Figure S3B). At its extremes, a learning rate closer to 1 means learning from only the most recent trials and a learning rate closer to 0 means learning from many previous trials. In this task, higher learning rates are adaptive, and correspond with faster block transitions and better performance. The inverse temperature determines choice stochasticity, or how deterministically the model acts (Figure S3C). High values of inverse temperature correspond to more deterministic choice functions — the agent will opt to choose the image with a higher value, even if the difference is small. Small values of inverse temperature correspond to more random behavior — the agent will still choose the image with lower value with reasonable probability. In this task, there is a more complex correspondence between inverse temperature and performance. High values of inverse temperature correspond to behavior that better maximizes reward when the better option is known, but tends to perseverate at block transitions. In general, higher values of inverse temperature correspond with better performance. The side bias determines the model’s preference for a stimulus location, regardless of relative image values (Figure S3D). Nonzero values of side bias are strictly maladaptive in this task and correspond to poorer performance.

To better estimate model parameters, we adopted a hierarchical Bayesian strategy to fit the reinforcement learning model and obtain monkey- and session-level parameter estimates (Figure 6A-C, S4). We related these parameter estimates to performance to gain better insight. We found that the learning rate improved with increased performance (Figure 6D; linear slope 2.47, *t*_7_ = 7.70, p < 0.0001). In contrast, the inverse temperature showed no significant linear association with performance (Figure 6E; linear slope −3.23, *t*_7_ = −2.00, p = 0.086). Because changes in learning rates and inverse temperatures can partially compensate for one another (small increase in learning rate can be compensated for by a small decrease in inverse temperature; (Daw et al., 2011)), we sought to measure their combined effect on P(choice). The maximal trial-by-trial change in P(choice), which partially accounts for this interaction, showed an increase with performance (Figure 6F; linear slope 1.00, *t*_7_ = 5.99, p = 5.5×10^−4^). Finally, the absolute value of side bias showed no change with performance (Figure 6G; linear slope −5.91, *t*_7_ = −1.80, p = 0.11; see Discussion).

**Figure 6:**
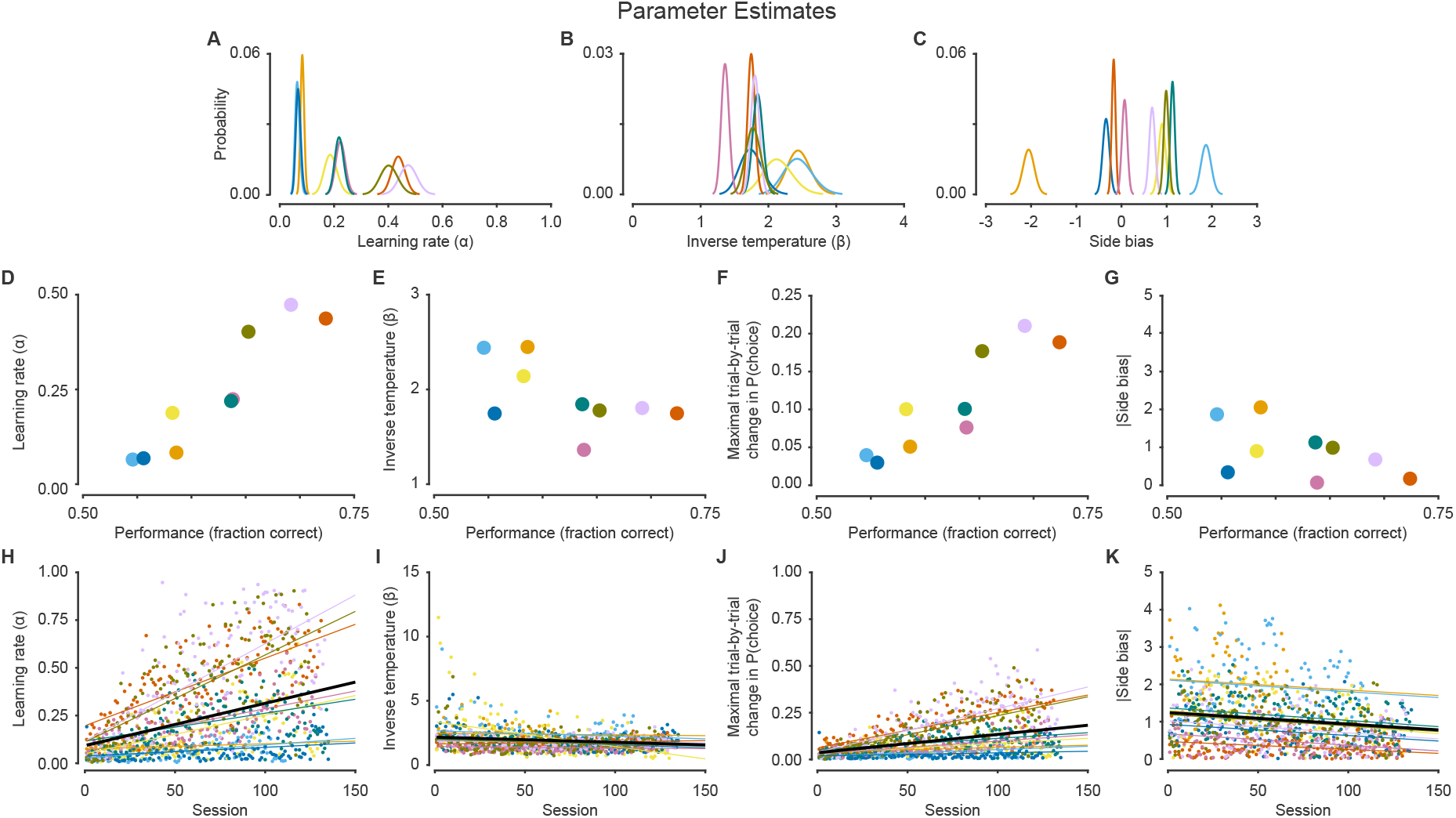
Relationship between model parameters, performance, and training. (A) Estimate learning rates for all monkeys. (B) Estimated inverse temperatures for all monkeys. (C) Estimated side biases for all monkeys. (D) As performance improved, the learning rate increased. (E) Inverse temperature showed no significant linear association with performance. (F) The maximal trial-by-trial change in P(choice), which partially accounts for the interaction of both the learning rate and the inverse temperature, increased as performance improved. (G) The absolute side bias showed no significant relationship with performance. (H) Learning rates improved with training. (I) The inverse temperature did not change throughout training. (J) The maximal trial-by-trial change in P(choice) increased with training. (K) Side bias decreased with training. Black line shows the fixed effect and thin colored lines show individual monkey random effects. Colors denote individual monkeys and are consistent between figures.

We next sought to estimate how these parameters changed with training. Learning rates increased with training, which yields better performance (Figure 6H; linear slope 2.22 × 10^−3^, *t*_1091_ = 3.78, p = 1.7 × 10^−4^). In contrast, inverse temperature showed no significant change with training (Figure 6I; linear slope −4.02 × 10^−3^, *t*_1091_ = −1.86, p = 0.06). This is noteworthy since animals consistently adopted suboptimal inverse temperatures and would have benefited from increased *β* values (Figure S4; see Discussion). The maximal trial-by-trial change in P(choice) improved with training (Figure 6J; linear slope 9.79 × 10^−4^, *t*_1091_ = 3.53, p = 4.7 × 10^−4^). Finally, side bias decreased with training, which permitted better performance (Figure 6K; linear slope −3.10 × 10^−3^, *t*_1091_ = −5.62, p < 0.0001). We obtained similar results after within-animal normalization by *z*-scoring, though with a small decrease in inverse temperature with training (Figure S6).

In summary, the reinforcement modeling approach provides a simple and compelling model for understanding how squirrel monkeys solve a reversal learning task and provides a tool for interpreting how the inner mechanisms relate to performance and how they change with training.

## Discussion

The power of comparative neuroscience to dissect cognition relies on the use of behavioral tasks that engage similar cognitive mechanisms in different species. Here, we show that squirrel monkeys solve a reversal learning task, a frequently-used behavioral paradigm, similarly to other species. We found that these animals integrate reward history over many trials to dictate choices, a commonly-observed reinforcement learning motif across species.

Using generative modeling, we explicitly tested a number of hypotheses about the strategies animals applied to harvest reward. We found that animal behavior was consistent with a remarkably simple strategy: reward history integration over many trials (~ 5-10) and a bias for a particular side. This model outperformed a number of other reasonable hypotheses. In particular, we found that a one learning rate model outperformed models with two learning rates. This is notable since models with two learning rates, which allow for separate learning from positive and negative reward prediction errors, are commonly found to better explain behavioral data (Frank et al., 2004; Grossman et al., 2020; Taswell et al., 2018; Averbeck, 2017; Dorfman et al., 2019; Gershman, 2015; Niv et al., 2012), although these tasks often have different reward statistics than what we used here.

We found that animals did not implement a pure or noisy win-stay/lose-shift strategy, either in isolation or mixed with a reinforcement learning strategy. Why didn’t animals approximate the optimal strategy in this task? Although win-stay/lose-shift is optimal on this particular task variant, it may not be adaptive across task variants in general. Reward probabilities vary drastically and dynamically in natural environments and, presumably, by integrating reward history, animals would continue to perform well if reward probabilities changed. Reward could be optimized by tweaking parameters (e.g. adjusting learning rates), rather than changing the entire behavioral strategy (Doya, 2002). Another potential reason is that incorrect choices still yielded reward, allowing animals to perform well enough despite using a suboptimal strategy.

Monkeys did not adopt optimal combinations of learning rates and inverse temperatures to maximize reward (Figure S5). Although animals had some exposure to the basics of the task (data not included), early in training, we would not expect animals to implement optimal parameter combinations. With training, they may approximate ideal parameter combinations with greater knowledge of task statistics. Indeed, we found that learning rates increased with training, which allows for better performance. Inverse temperatures, however, did not increase, which would be expected to optimize reward. Lower inverse temperatures meant monkeys made choices more randomly. This finding is consistent with the notion that monkeys maintained a high level of exploratory behavior, which may be a ubiquitous feature of behavior even when task demands encourage more deterministic choice behavior (Pisupati et al., 2021; Ebitz et al., 2019). Limited attention may also contribute to suboptimal exploratory behavior.

Side-specific biases were a key feature of behavior in this task. These low-level idiosyncratic tendencies are often ubiquitous features of animal behavior. These side biases are likely not controlled by the same brain regions that engender flexible behavior (Balleine and O’doherty, 2010; Bari et al., 2019). The reduction in side bias and increase in learning rate were correlated during training (Figure 6H-K), likely because both processes yield an improvement in performance, but would likely be independent following neural manipulation (Bari et al., 2019).

One potentially inconsistent finding is that the side bias metric decreased with performance (Figure 3C, 5C), while the bias parameter from the reinforcement learning model did not show a statistically-significant change (Figure 6G). This is likely because this particular analysis was underpowered. When we analyzed data at the session level, we observed a significant decrease in both the mean and variance of the bias parameter (data not shown). For poor performance, side bias was highly variable. This is because poor performance could be the result of a strong bias to one side, or it could be the result of random, reward-insensitive behavior with no side bias. With better performance, side bias decreased in mean and variance, because a strong bias places an upper bound on performance, no matter how reward-sensitive animals are.

Generative modeling allows us to test hypotheses that may be beyond the reach of simple summary statistics. For example, it is reasonable that animals could have computed action values in addition to a side bias (e.g. in a task variant where computing action values may be adaptive). It’s not clear how the choice-based win-stay/lose-shift analyses we used (Figure 2H), which can test whether animals implement reward sensitivity to choices vs side bias, would help if animals implemented a mixture of the two. With the generative modeling approach, as long as the model is recoverable, then this hypothesis would be simple to test (Wilson and Collins, 2019).

Our modeling approach, while generally successful, did not perfectly recapitulate all behavioral features. One notable failure was the inability to capture the slight recovery of performance in the one trial immediately after a Reversal block began (compare Figures 2C, 4C). Interestingly, we found that simulated data with a mixture of reinforcement learning and win-stay/lose-shift was able to partially recapitulate this phenotype. However, the fact that none of these models fit animal behavior well (Table S1) argues that win-stay/lose-shift is not a cardinal feature of behavior, at least given our model selection pipeline. Interestingly, win-stay/lose-shift may only be a strategy animals implement on particular trials (Iigaya et al., 2017), which may disfavor a model that assumes win-stay/lose-shift is implemented on every trial. Perhaps squirrel monkeys implement win-stay/lose-shift only following large magnitude negative reward prediction errors, accounting for behavior in the trials immediately after a block change, and otherwise implement reinforcement learning. Learning rates might also change as a function of recent reward statistics, yielding non-stationary behavioral strategies (Behrens et al., 2007; Nassar et al., 2012; Grossman et al., 2020).

One strength of generative modeling is that it allows for interpretable insights into manipulations, particularly across species. Parameter estimates (Figure S4) may be compared across groups to gain insight into the effects of disease or manipulations (Kanen et al., 2021; Aylward et al., 2019; Huys et al., 2013). A complementary approach is to extract the latent variables governed by these parameters and correlate them with neural activity (Samejima et al., 2005; Bari et al., 2019; Findling et al., 2019). Insights at the level of parameters or latent variables may aid the development of novel therapies, since the development pipeline for nervous system therapeutics often stalls due to lack of objective biomarkers of success (Kola, 2008; Paulus et al., 2016). Since these types of models have theoretical underpinnings, parameter changes may be interpreted through the lens of theory. For example, the volatility of the environment should modulate learning rates (Behrens et al., 2007), beliefs about the causal structure of the environment should modulate asymmetric learning from good and bad outcomes (Dorfman et al., 2019), and the complexity of action space should govern the inverse temperature and perseveration (Gershman, 2020).

The reinforcement learning model we chose is a fairly general algorithm that is not specific to the task the monkeys performed. In fact, to best study the cognitive mechanisms underlying this algorithm across species, we may need to adjust the task across species to account for species-specific differences. Humans performing a deterministic reversal learning task would almost certainly discover that win-stay/lose-shift was the optimal policy and exploit it. Rhesus macaques overtrained on a deterministic reversal learning paradigm eventually learn expected block lengths (Jang et al., 2015). Therefore, to best study this algorithm in humans and macaques may require a probabilistic reversal learning task without overtraining (to avoid win-stay/lose-shift policies), or a task where the probabilities drift slowly across time, without clear reversals (to avoid learning expected block lengths; (Daw et al., 2006)).

The behavior of squirrel monkeys is notable for the larger number of within-session reversals compared to rodents and marmosets, which is more comparable to macaques and humans (Izquierdo et al., 2017). This means that well-trained squirrel monkeys may more readily approximate human strategies, which would significantly aid the ability to translate insights. Our results highlight the utility of reinforcement learning modeling and validates squirrel monkeys as a useful behavioral neuroscience model.

## Acknowledgements

We thank Samantha O. Brown, and Rodrigo A. Montoro for assistance with monkey training. We thank Frederick Barrett, Matthew Johnson, Sandeep Nayak, Manoj Doss, Justin Strickland, Ceyda Sayalı, and Cooper Grossman for valuable discussions at various stages of the project. We thank Cooper Grossman for providing feedback on an earlier version of the manuscript.

---

**Figure S1:**
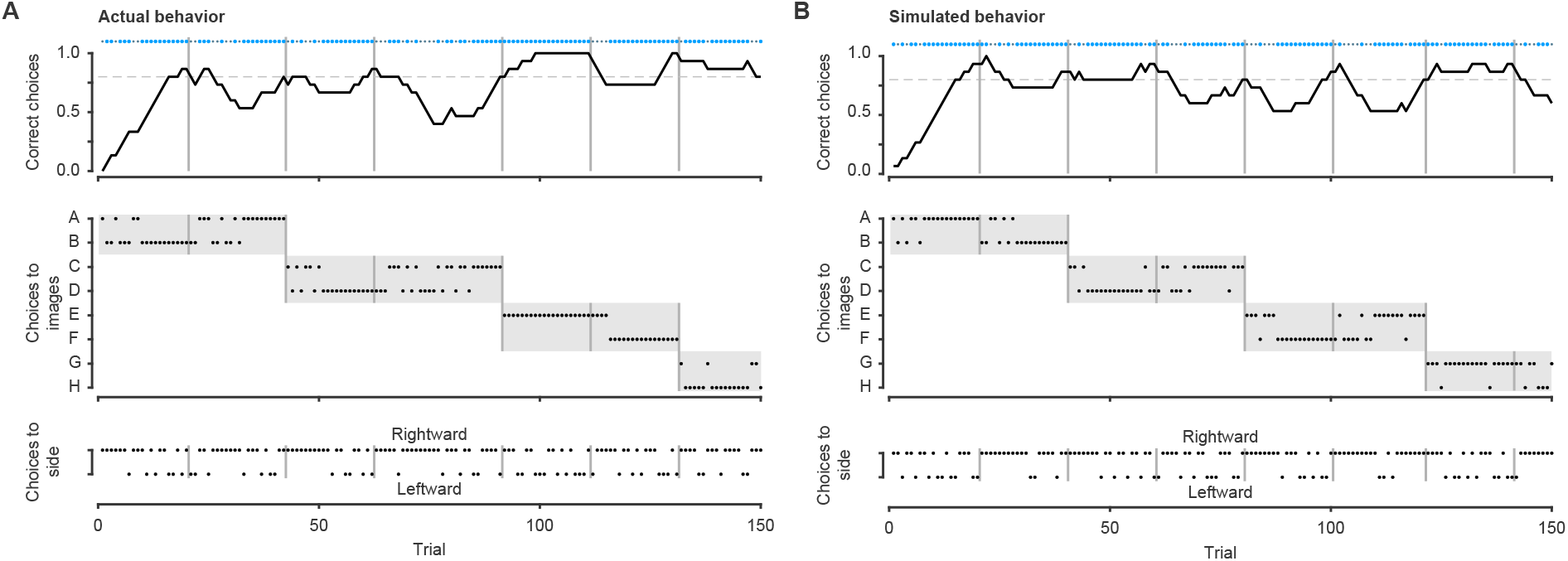
Raw behavior for actual and simulated sessions. (A) Top panel: Correct choices as a function of trial. Large dots indicate a big reward choice and small dots indicate a small reward choice. The black line shows the fraction correct in the past 15 trials. The dashed line is the performance threshold (80% correct) used to trigger block transitions. Vertical grey lines indicate block transitions. Middle panel: Choices to images as a function of trial in the same format as 1C. Black dots indicate a choice to a respective image. Bottom: Choices to a side as a function of trial. Rightward (leftward) choices are indicated with a black dot on the top (bottom) of the figure. This session demonstrates a slight rightward side bias. (B) Behavior from the same session was fit to the reinforcement learning model to estimate parameters. These parameters were used to simulate an entirely new, synthetic data session.

**Figure S2:**
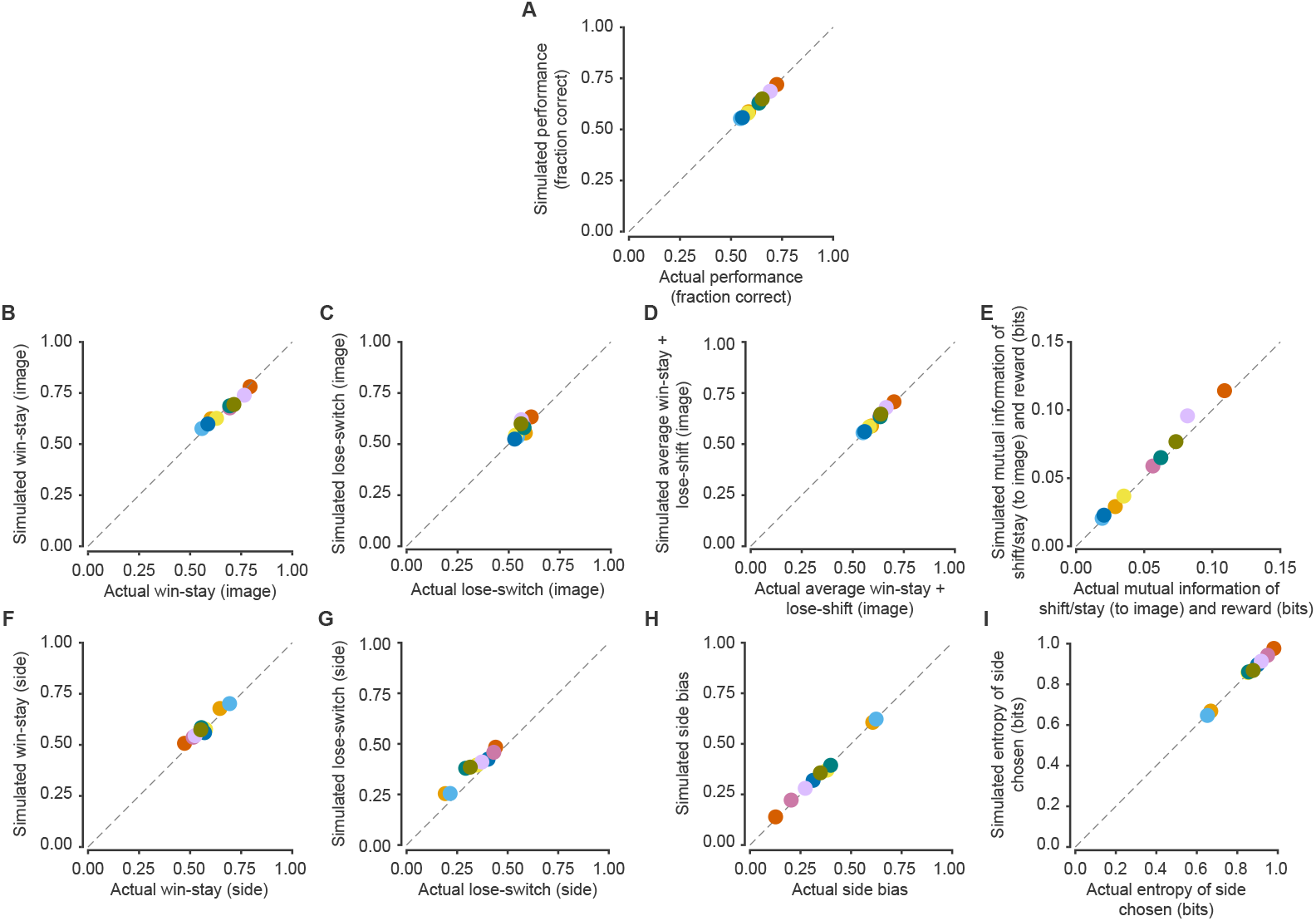
Comparison of behavioral features for actual and simulated behavior. To compare how well actual and simulated behavioral features match, we compute the mean difference between actual and simulated behavioral features [Mean (actual minus simulated) 95% CI] for each panel. (A) Average performance for each session. [−0.0015 – 0.0042] (B) Image-based win-stay [−0.0070 – 0.0130] (C) Image-based lose-shift [−0.0303 – 0.0015] (D) Image-based average win-stay + lose-shift [−0.0043 – 0.0064] (E) Mutual information of stay/shift and reward on the previous trial [−0.0082 – −0.0021] (F) Side-based win-stay [−0.0265 – −0.0062] (G) Side-based lose-shift [−0.0632 – −0.0367] (H) Side bias [−0.0097 – 0.0015] (I) Entropy of side chosen distribution [−0.0012 – 0.0052]. Colors denote individual monkeys and are consistent between figures.

**Figure S3:**
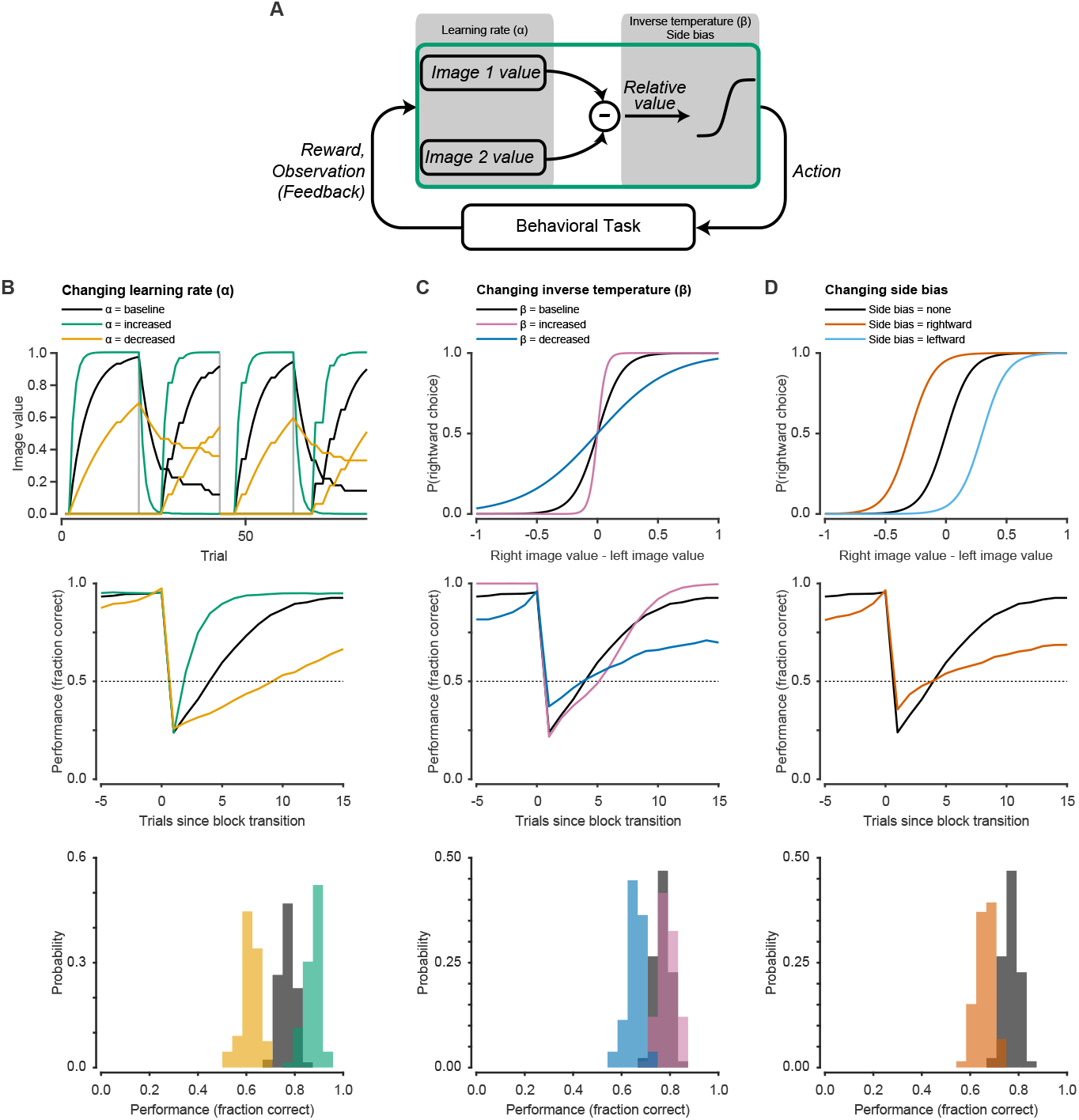
Model illustration and effects of varying parameters. (A) Illustration of reinforcement learning model. Image values are updated by feedback via reward prediction errors (the discrepancy between predicted and actual rewards). This process is governed by the learning rate (*α*). The relative image value is mapped through a softmax function to produce a choice. This process is governed by the inverse temperature (*β*) and a side bias parameter. (B) Increasing the learning rate results in faster accumulation of reward value information. This results in faster block transitions and better overall performance. Decreasing the learning rate has the opposite effect. (C) Increasing the inverse temperature results in more deterministic choice behavior. Decreasing the inverse temperature makes choices more random. In this example, increasing the inverse temperature results in slower block transitions but more deterministic behavior after enough trials have elapsed, resulting in improved performance. Decreasing the inverse temperature has the opposite effect. Unlike the learning rate, the optimal inverse temperature is not at an extreme value but depends on the trials to criterion. Greater trials to criterion will favor a larger *β*. (D) Side bias results in increased choices of one particular side. Side bias is purely maladaptive and results in poorer overall performance.

**Figure S4:**
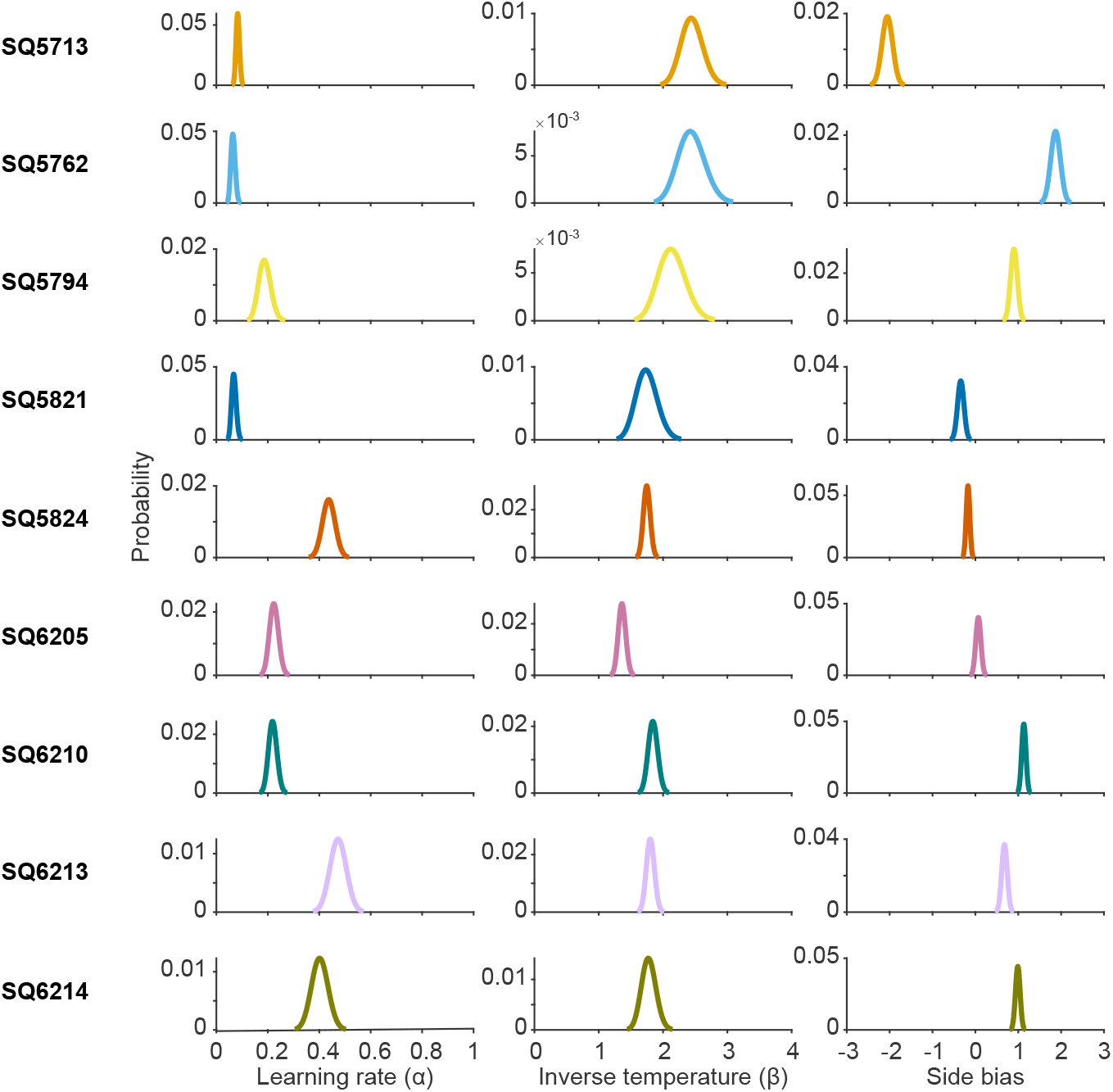
Parameter estimates for each monkey. Estimated learning rates, inverse temperatures, and side biases for all monkeys included in this study. Colors indicate the color used for that monkey throughout figures.

**Figure S5:**
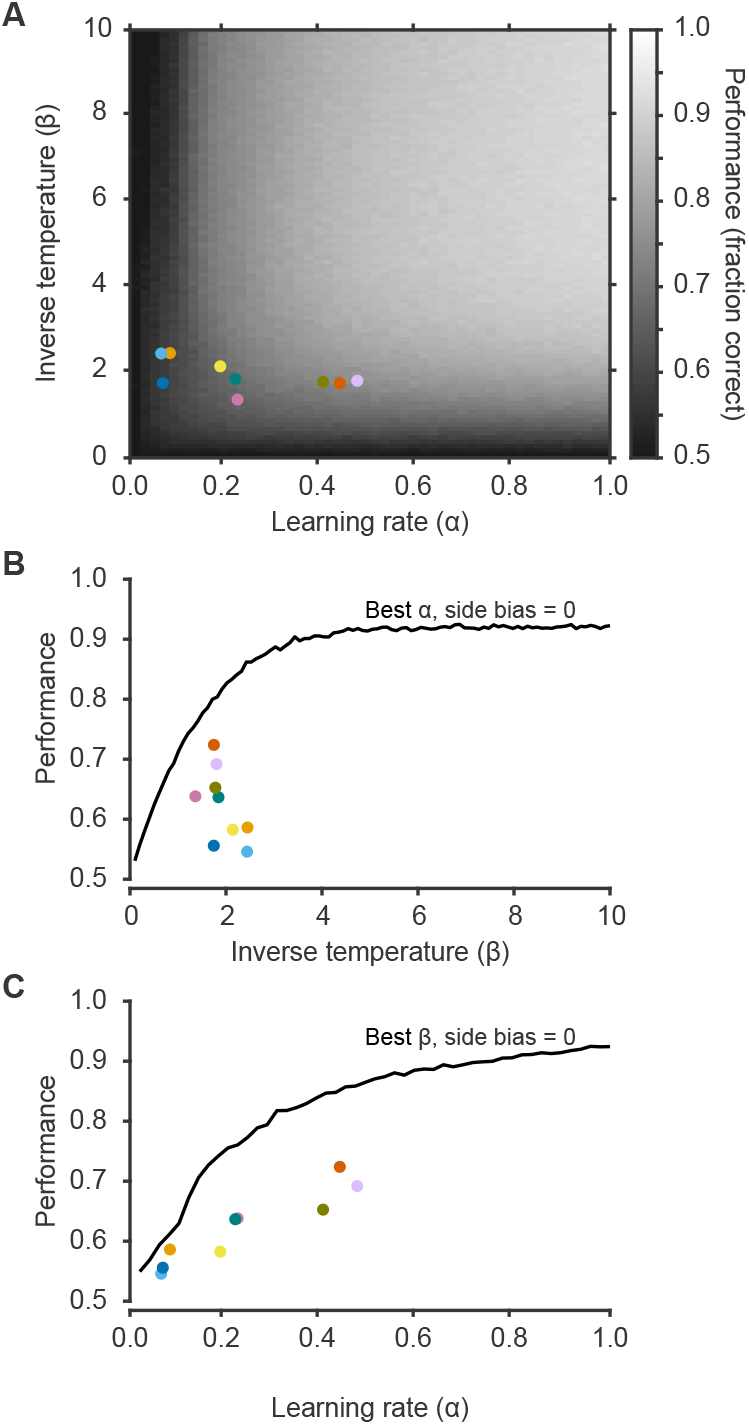
Performance as a function of learning rate (*α*) and inverse temperature (*β*). Each simulation was run for 66 sessions, each 2000 trials long, over 50 *α* values, 100 *β* values, and side bias fixed at 0. (A) Heatmap of performance for combinations of learning rates and inverse temperatures, with side bias fixed at 0. Performance is poor at low learning rates (regardless of the inverse temperature) and low inverse temperatures (regardless of the learning rate). In general, there is a large range of learning rates and inverse temperatures that permits adaptive behavior. Individual monkeys are shown with colored dots. Monkeys consistently maintain a suboptimal *α*/*β* combination. (B) Performance as a function of inverse temperature for the best learning rate and side bias = 0. Optimal performance is achieved at *β* ≳ 4. (C) Performance as a function of learning rate for the best inverse temperature and side bias = 0. Optimal performance is achieved at *α* = 1. Colors denote individual monkeys and are consistent between figures.

**Figure S6:**
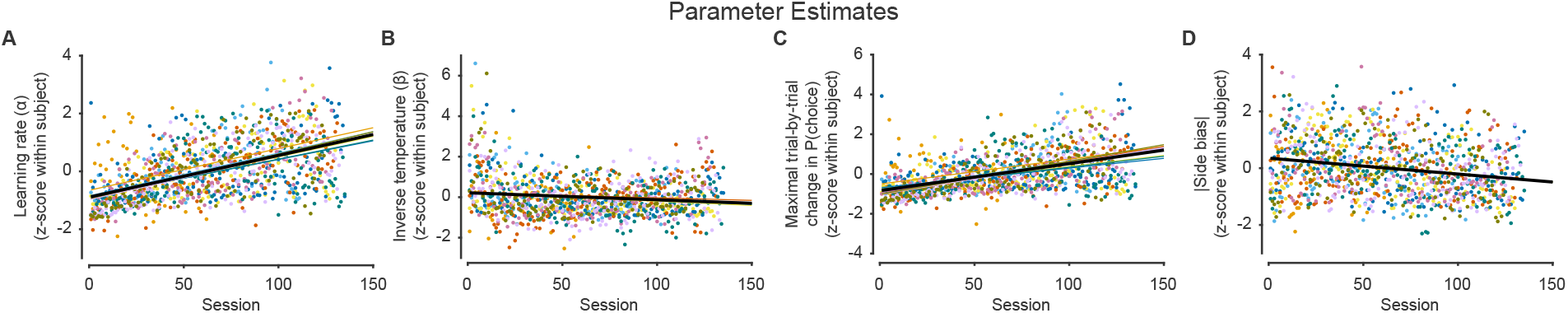
Within-subject normalization of parameters results in similar changes with training. (A) Normalized learning rates improved with training (linear slope 1.44 × 10^−2^, *t*_1091_ = 17.33, p < 0.0001). (B) Normalized inverse temperatures decreased with training (linear slope −3.46 × 10^−3^, *t*_1091_ = −4.05, p < 0.0001). (C) Normalized maximal change in P(choice) increased with training (linear slope 1.36 × 10^−2^, *t*_1091_ = 11.13, p < 0.0001). (D) Normalized absolute side bias decreased throughout training (linear slope −5.61 × 10^−3^, *t*_1091_ = −7.11, p < 0.0001). Colors denote individual monkeys and are consistent between figures.

**Table S1:** Model comparison. Comparison of negative log likelihood (LH) and Bayesian information criterion (BIC) values for the four models best fit to at least one monkey, and one noise model. The four best-fit models are variations of the Rescorla-Wagner (RW) model with a side bias. The noise model is a side bias only model. LH and BIC values are sums across all sessions for individual monkeys. Colors in the Monkey column indicate the color used for that monkey throughout figures. Gray highlights the best model (smallest BIC) for each monkey.

| Monkey | Model |  |  |  |  |  |  |  |  |  |
| --- | --- | --- | --- | --- | --- | --- | --- | --- | --- | --- |
|  | RW +<br>Side bias<br>(3 parameters) |  | RW +<br>Side bias +<br>Reversal<br>mechanism<br>(3 parameters) |  | RW +<br>Side bias +<br>Forget unchosen<br>(4 parameters) |  | RW +<br>Side bias +<br>Forget unchosen +<br>reward coded as<br>[0.25 1]<br>(4 parameters) |  | Side bias<br>(1 parameter) |  |
|  | LH | BIC | LH | BIC | LH | BIC | LH | BIC | LH | BIC |
| SQ5713 | 3685 | 8363 | 3756 | 8505 | 3639 | 8601 | 3638 | 8598 | 4586 | 9503 |
| SQ5762 | 6271 | 14198 | 6366 | 14389 | 6152 | 14513 | 6114 | 14438 | 7288 | 15129 |
| SQ5794 | 9139 | 20186 | 9239 | 20385 | 8833 | 20211 | 8791 | 20127 | 10595 | 21826 |
| SQ5821 | 11387 | 24796 | 11512 | 25046 | 11310 | 25315 | 11272 | 25240 | 12446 | 25566 |
| SQ5824 | 9588 | 21135 | 9522 | 21003 | 9260 | 21132 | 9285 | 21181 | 13089 | 26831 |
| SQ6205 | 10776 | 23475 | 10801 | 23525 | 10583 | 23728 | 10585 | 23732 | 12573 | 25786 |
| SQ6210 | 9483 | 20988 | 9559 | 21140 | 9199 | 21093 | 9223 | 21141 | 11835 | 24344 |
| SQ6213 | 9341 | 20675 | 9313 | 20619 | 8750 | 20157 | 8777 | 20212 | 12556 | 25777 |
| SQ6214 | 8535 | 18925 | 8591 | 19037 | 8091 | 18655 | 8085 | 18645 | 11073 | 22764 |

